# Diversity and varying predation capacities of Amoebozoa against opportunistic vibrios in contrasting Mediterranean coastal environments

**DOI:** 10.1101/2025.03.14.643055

**Authors:** Etienne Robino, Angélique Perret, Cyril Noel, Philippe Haffner, Laurent Intertaglia, Marion Richard, Noémie Descamps, Axelle Sellier, Laura Onillon, Philippe Lebaron, Delphine Destoumieux-Garzón, Guillaume M. Charrière

## Abstract

Free-living amoebae (FLA) are ubiquitous and can be found in many types of environments including soil, freshwater, air and marine environments. They feed on various microorganisms and can play an important role in the food web and its dynamics. We previously described that FLAs belonging to the *Vannella* genus from the oyster farming area of the Thau lagoon in France could establish stable interactions with *Vibrionaceae* and could play a role in the selection of some virulence factors, potentially influencing pathogen dynamics. To further investigate the ecological interactions between FLA populations and Vibrionaceae in Mediterranean coastal waters, we conducted monthly sampling for one year at three contrasting sites. Free-living amoebae populations were isolated by culturing water and sediment samples on different bacterial lawns, including *E. coli* or *V. tasmaniensis*, *V. crassostreae* and *V. harveyi*. Amoebozoan diversity was analyzed using v4-18S barcoding and revealed distinct communities between the sediment and the water column, with Vannellidae significantly enriched in the water column whereas Paramoebidae were significantly enriched in the sediments. Selection of grazers on different bacterial lawns revealed that *V. tasmaniensis* inhibited the growth of most Vannellidae, whereas *V. crassostreae* inhibited the growth of a large proportion of Paramoebidae. These differences were further confirmed in functional grazing assays using isolates belonging to each Amoebozoa taxonomic group. Overall, our results highlight the need for more comprehensive studies of the diversity and population dynamics of Amoebozoa in marine environments, and that the role of these diversified grazers in shaping vibrio communities is complex and still poorly characterized.

## Introduction

Grazing protists, such as free-living amoebae (FLA), are found in many different environments. Free-living amoebae are mainly found in freshwater and marine environments, but also in soil and air, and can be associated with various hosts (Rodríguez-Zaragoza, 1994; Samba-Louaka et al., 2019). The diversity of FLAs has been studied more extensively in freshwater environments than in marine environments, especially because human pathogenic amoebae are mostly found in freshwater (Visvesvara et al., 2007). Amoebae are a polyphyletic group that branches along the eukaryotic tree and belongs to four supergroups, Amoebozoa, Excavata, Opisthokonta and SAR. Amoebozoa is the only group that contains only FLAs (Lahr et al., 2011). Free-living amoeba feed on various microorganisms and digest them by phagocytosis. They can feed on bacteria, yeasts, fungi, algae or other protists (De Moraes and Alfieri, 2008; Radosa et al., 2019; Salt, 1968; Smirnov et al., 2011). As heterotrophic protists, they are central members of the food web in the environment, and their predation on bacteria is a major factor shaping microbial communities (Batani et al., 2016; Corno and Jürgens, 2008; Gao et al., 2019; Jürgens et al., 1999; Pernthaler, 2005). Protist dynamics (e.g., abundance and diversity) are influenced by many environmental factors such as prey type and abiotic factors such as salinity, temperature, and oxygen availability (Amaro et al., 2015; Guillou et al., 2001; Kim et al., 2014; Orsi et al., 2011; Smirnov, 2007). We observed previously that the diversity of FLA in the Mediterranean Thau lagoon near the oyster farming area is relatively low, with amoebae mainly belonging to the *Vannella* genus, and some of these *Vannella* could establish stable interactions with Vibrionaceae (Robino et al., 2020). Vibrios are γ-proteobacteria that live in aquatic environments, both freshwater and marine. They are ubiquitous and associated with many hosts from protozoa to metazoans, and these associations range from symbiosis to pathogenesis (Mandel and Dunn, 2016; Takemura et al., 2014). Vibrios are involved in various vibriosis in marine animals and can have devastating impacts in aquaculture, including oyster farming. (Dubert et al., 2017; Le Roux et al., 2016). For example, they can be involved in mortality events of juvenile oysters *Crassostrea gigas* (De Lorgeril et al., 2018). Immunosuppressed oysters after OsHV-1 infection are colonized by opportunistic vibrios that cause septicemia, leading to animal death. These opportunistic vibrios, which have been implicated in Pacific oyster syndrome mortality, belong to *V. crassostreae*, *V. tasmaniensis* and *V. harveyi* species (Gay et al., 2004; Lemire et al., 2015; Oyanedel et al., 2023). Interestingly, cytotoxicity against hemocytes by *V. tasmaniensis* LGP32, *V. crassostreae* J2-9, and *V. harveyi* A01 has been shown to be a major driver of their virulence, but relies on species-specific mechanisms (Oyanedel et al., 2023; Rubio et al., 2019). The acquisition of such anti-eucaryotic activities can result from the long co-evolutionary history between bacteria and FLAs. Some bacteria have evolved resistance to amoeba predation through extracellular and intracellular defense mechanisms that play a role in virulence against animal hosts (Matz and Kjelleberg, n.d.; Pernthaler, 2005). Therefore, amoebae are considered as evolutionary ancestors of interactions and act as a training ground for intracellular pathogenic bacteria by favoring the selection of virulence factors through coincident evolution (Diard and Hardt, 2017; Molmeret et al., 2005). In the case of vibrios, *Vibrio cholerae* has been shown to be able to survive in *Acanthamoeba castellanii* amoebae and to use some virulence genes that play a minor role in the interaction with the incidental human host (Van Der Henst et al., 2018). However, when *Vibrio cholerae* leaves *Acanthamoeba castellanii* by exocytosis of the vibrio-containing food vacuole, this can increase its virulence against mammalian hosts (Espinoza-Vergara et al., 2019). In the case of *V. vulnificus*, the virulence factor MARTX type III was shown to be involved in resistance to grazing by *Neoparamoeba permaquidensis* isolated from fish gills (Lee et al., 2013). By comparative cell biology, we have previously shown that the oyster opportunistic pathogen *V. tasmaniensis* LGP32 is able to resist phagocytosis by the environmental marine amoeba *Vannella* sp. AP1411 using some virulence factors also involved in oyster pathogenesis, in particular the secreted metalloprotease Vsm and the copper efflux pump CopA (Robino et al., 2020). Finally, lipopolysaccharide O-antigen variations in *Vibrio splendidus* strains were shown to determine resistance to grazing by *Vannella* sp. AP1411 (Oyanedel et al., 2020). Taken together, these different reports suggest that amoeba-vibrio interactions are diverse and need to be further studied to better evaluate their role in vibrio dynamics and pathogen emergence.

To study the ecology and interactions between FLA populations and *Vibrionaceae* in Mediterranean coastal waters, in the present report we conducted monthly sampling for one year in three contrasting environments. In the Thau lagoon, used for oyster farming and where oyster mortality due to POMS occurs annually (Richard et al., 2021), but also in the Mediterranean Sea outside the port of Sète, and near the marine protected area of Banyuls-sur-Mer. Free-living amoebae populations were isolated by culturing water and sediment samples on different bacterial lawns, including *E. coli* SBS363, *V. tasmaniensis* LGP32, *V. crassostreae* J2-9 and *V. harveyi A01*. Using 18S barcoding, we found that Amoebozoa dominated the FLA populations with higher diversity in the sediments than in the water column, with distinct communities and site-specific diversity in the sediments. One of the most striking differences was that Vannellidae are abundant and specifically enriched in the water column, whereas Paramoebidae are abundant and specifically enriched in the sediments. Furthermore, we show that these two families of amoebozoans have contrasting predation capacities against two different species of vibrios of the Splendidus clade *V. tasmaniensis* LGP32 or *V. crassostrae* J2-9, which was further confirmed functionally using amoeba isolates belonging to each family. Overall, our results highlight that the diversity of Amaoeboza is still understudied and needs to be further characterized in different marine environments, and that the nature of amoeba-vibrio interactions may occur at very different taxonomic levels and can play a role in the dynamics of opportunistic pathogens in coastal marine waters.

## Materials and methods

### Bacterial strains, amoeba strain and growth conditions

*E. coli* strain SBS363 was grown in Luria-Bertani (LB) or LB-agar (LBA) at 37°C for 24 hours prior to experiments. *V. tasmaniensis* LGP32, *V. crassostreae* J2-9, and *V. harveyi* A01 were grown in LB + NaCl 0.5M or LBA + NaCl 0.5M at 20°C for 24 hours prior to experiments. Vibrios strains carrying the pMRB-GFP plasmid were grown in LB + NaCl 0.5M supplemented with chloramphenicol (10µg.mL^-1^) at 20°C for 24 hours prior to experiments. *Vannella sp*. AP1411 (isolated in a previous study, (Robino et al., 2020) and *Paramoeba atlantica* strain CCAP1560/9 (purchased from the CCAP collection, Scotland, UK) were cultured at 18-20°C in 3 ml of 70% sterile seawater (SSW) supplemented with 200 µL of a *Escherichia coli* SBS363 suspension (OD600=20).

### Sampling

Sampling was performed monthly for one year between 2017 and 2018 in three contrasting environments in southern France. Water and sediment were collected in the Thau lagoon adjacent to oyster beds at the Bouzigues station Ifremer-REPHY (GPS: N 43°26’.058’ - E 03°39’.878’), in the open sea near the port of Sète (GPS: N 43°23’.539’ - E 03°41’.933’), and near the Banyuls-sur-Mer marine reserve at the SOLA station (GPS: N 42°29’300’ - E 03°08’700’) (Figure S1). The surface water (1 meter depth) was sampled with a hydrobios bottle at the three sites, and the first centimeters of sediments were sampled with cores taken by divers at 9, 10 and 30 meters depth respectively at Thau lagoon, Sète and Banyuls-sur-Mer. The sediments from Thau and Sète were muddy and sandy, respectively, while the sediments from Banyuls-sur-Mer had a mixed composition. Water was filtered on the boat using a 63 µm pore size nylon filter and then refiltered in the laboratory using a 5 µm pore size MF-Millipore membrane. The 5 µm pore size membrane was then cut into four pieces and three quarters were placed upside down on a lawn of *E. coli* SBS363 seeded on 70% SSW agar, while the other whole filter was cryopreserved at -80°C. One gram of sediment was placed in the center of a lawn of *E. coli* SBS363 seeded on 70% SSW agar in triplicate and incubated at 20°C while the same type of samples were cryopreserved at minus 80°C. Briefly, after 2 weeks, the amoebae were rinsed with 70% SSW and one milliliter of the rinsed solution was cryopreserved at -80°C for each condition. The same steps were performed with *V. tasmaniensis* LGP32, *V. crassostreae* J2-9, and *V. harveyi* A01 in May and October 2017 during oyster mortality events and in January and February without mortality events. A total of 76 samples from *E. coli* plates and 84 samples from vibrios plates were collected for diversity analysis using 18S barcoding.

### 18S barcoding data processing

Total DNA from rinsed agar plates was extracted using a Blood and Tissue kit from Marcherey Nagel with an adapted initial lysis phase using 0.1mm zirconium beads under string agitation in T1 buffer. Eukaryote diversity was analyzed by barcoding based on the polymorphism of the v4 loop of the 18S rRNA coding gene. PCR were performed using universal primers at an annealing temperature of 53°C (TAReuk454FWD-illumina: 5’-TCG TCG GCA GCG TCA GAT GTG TAT AAG AGA CAG YRC CAG CAS CYG CGG TAA TTCC-3’ and TAReukRev3 Illumina: 5’-GTC TCG TGG GCT CGG AGA TGT GTA TAA GAG ACA GYR ACT TTC GTT CTT GAT YRA-3’) (Stoeck et al., 2010). Paired-end sequencing (300 bp read length) was performed using the GenSeq platform (Labex CEMEB, Montpellier, France) on the MiSeq system (Illumina). Data is available under the BioProject PRJEB87851 on ENA database https://www.ebi.ac.uk/. Raw data were then processed using the SAMBA v3.0.1 pipeline (https://gitlab.ifremer.fr/bioinfo/workflows/samba), a standardized and automated workflow for meta-barcoding analyses. This workflow, developed by SeBiMER (Ifremer’s Bioinformatics Core Facility), is an open source modular workflow for processing eDNA metabarcoding data. SAMBA, developed by SeBiMER (Ifremer’s Bioinformatics Core Facility), is an open-source modular workflow for processing eDNA metabarcoding data. SAMBA was developed using the Nextflow workflow manager (Di Tommaso et al., 2017) and allows the execution of three main components: data integrity verification, bioinformatics processes, and statistical analyses.

### Amoebozoa diversity analysis on different nutritive sources

Seventy two samples were obtained from *E. coli* agar plates, 6 samples that yielded less than 1000 sequences were removed (10-17-Ba-S-E-coli, 11-17-Th-S-E-coli, 06-17-Th-S-E-coli, 09-17-Th-S-E-coli, 12-17-Ba-S-E-coli, and 11-17-Ba-S-E-coli) and 2 samples were considered outliers due to a species composition that was completely different from the rest of the samples (08-17-Th-W-E-coli and 09-17-Ba-W-E-coli). Analysis of the remaining 64 samples resulted in a total of 2,780,405 sequences after data integrity checks. QIIME 2 was used for primer removal using the Cutadapt plugin, sequence quality check was performed using the DADA2 R package (removal of low-quality sequences, assembly of forward and reverse sequences, and removal of chimeras), and ASV clustering was done using the dbOTU3 algorithm to obtain a total of 2556 ASVs (Bolyen et al., 2019; Callahan et al., 2016; Hedjazi et al., 2011; Olesen et al., 2017). PR2 database was used for taxonomic assignment (version 4.13.0) with taxonomy filtering to select only ASVs belonging to the phylum of Amoebozoa, which consists only of FLAs (Guillou et al., 2012). A total of 465 ASVs were regrouped into a table containing the names of the ASVs with their taxonomic affiliations, as well as the names of the samples containing the ASVs and number of reads identified for each sample (Table S2). For the analysis of the samples acquired on the four different nutrient sources (*E. coli* SBS363, *V. harveyi*A01, *V. tasmaniensis* LGP32 and *V. crassostreae* J2-9), 112 samples were obtained (i.e 84 samples from Vibrios lawn and 28 samples from *E. coli* lawn), 6 samples with less than 900 sequences were removed (02-18-Ba-S-LGP32, 01-18-Th-S-A01, 10-17-Ba-S-E-coli, 05-17-Th-S-LGP32, 05-17-Ba-S-J2-9 and 05-17-Se-S-J2-9). Analysis of the 106 remaining samples resulted in a total of 5,298,844 sequences after data integrity checks. Through QIIME 2, we performed primer removal using the Cutadapt plugin, sequence quality check using the DADA2 R package, and ASV clustering using the dbOTU3 algorithm to obtain a total of 3084 ASVs (Bolyen et al., 2019; Callahan et al, 2016; Hedjazi et al., 2011; Olesen et al., 2017). Taxonomic affiliation was performed using the PR2 database and taxonomy filtering to select only ASVs belonging to the phylum Amoebozoa. We obtained a total of 474 ASVs, which were regrouped into a table containing the names of the ASVs with their taxonomic affiliations, as well as the names of the samples containing the ASVs and read counts identified for each sample (Table S5).

SAMBA then performed extensive analyses of alpha and beta diversity using custom R scripts (R Core Team, 2020). Alpha diversity was examined using the Chao1, Shannon, and InvSimpson indices. Beta diversity was investigated using Principal Coordinate Analysis (PCoA) with DEseq2 normalization method and Unifrac distance matrix. Repartition of the specifics and shared ASVs to the different conditions highlighted the taxonomy for each variable or combined variables and were plotted using UpsetR V1.4.0. The phylogenetic tree was performed using MAFFT and FastTree (maximum likelihood tree) and annotated using iTOL software, highlighting fraction, sites and season variables (Letunic and Bork, 2021).

### Grazing assay

To prepare the co-culture of vibrios and amoebae, 1 mL of Vibrio overnight culture (3.10^9^ bacteria. mL^-1^) was mixed with 100 µL of three day old *Vannella sp.* AP1411 or *Paramoeba atlantica* culture (5.10^5^ cells. mL^-1^) or with 100 µL of 70% SSW for the control condition. A volume of 50 µL per well of the mixed culture was seeded on 500 µL of 1% SSW agar in a 24-well plate with a transparent flat bottom. Amoebae and bacteria were carefully homogenized in the wells, allowed to dry by incubating for 4 hours at room temperature under a sterile hood, and then incubated at 18°C in a humidified atmosphere. GFP fluorescence intensity was measured daily for 7 days using a TECAN plate reader (λex 480 nm/λem 520 nm). To estimate the effect of amoeba grazing activity on the abundance of live GFP-expressing vibrios, the fluorescence intensity of wells containing amoebae was compared to the fluorescence of vibrios in the absence of amoebae and expressed as a ratio. Each condition was performed in triplicate and the results shown are representative of three independent experiments.

Error bars represent the standard error of the mean (±SEM). Statistical analysis was performed using RM-ANOVA on the independent experiments.

## Results

### Amoebozoa diversity varies greatly between the water column and sediments

The dynamics of grazers remains poorly described in Mediterranean coastal environments and even less in lagoons close to oyster farms. Therefore, we attempted to describe the diversity and distribution of cultivated grazers in Mediterranean coastal environments by studying 3 contrasting environments. Monthly sampling was carried out over an entire year in an environment close to the oyster culture tables in the Thau lagoon (Bouzigues), in an open sea site outside the port of Sète, and a last more distant and protected site in the Banyuls-sur-Mer area adjacent to the marine protected area (Figure S1). Sediments and water were sampled at the 3 sites every month for a whole year, from March 2017 to March 2018. To recover cultivable amoebae, samples were placed on non-nutritive plates with a layer of *E. coli* SBS363 as a non-pathogenic and permissive nutritive food source to isolate the highest diversity possible under these conditions.After 15 days of incubation at 20°C, total cultivable diversity was estimated by sequencing the v4 hypervariable region of 18S rRNA using universal primers. We then used the SAMBA pipeline to process the sequencing data (see materials and methods section). Due to the lack of taxonomic references in 18S SSU databases for some protist groups, we chose to focus on ASVs belonging to Amoebozoa, as they represent the majority of ASVs with a taxonomic affiliation and this taxonomic group contains only FLAs. First analyses showed that the alpha-diversity and beta-diversity of Amoebozoa differed significantly between the two different sampling fractions (e.g. sediment or water column), independent of sampling location (Figure 1A and B). The Chao1 index (estimator of species richness) was found to be significantly different, with a median Chao1 in the water column of 6.5 ASVs compared to a median of 18 ASVs for the sediments, indicating that alpha diversity tends to be lower in the water column than in the sediments (ANOVA test; p = 0.0138, Table S2). The other two alpha diversity indices, Shannon and InvSimpson, were not significantly different, indicating that the differences between the fractions were mostly due to low abundance species. For beta-diversity, PCoA revealed different Amoebozoa communities between water and sediment samples (Adonis test; Adonis R^2^ = 0.19, p = 0.0001, Table S3, Figure 1A). In addition, sediment samples appeared more grouped than water samples, suggesting a more stable community over the sampling months (Figure 1A). The repartition of ASVs grouped by fractions showed that of the 465 ASVs, 312 ASVs (67.1%) were found only in sediments, while 122 ASVs (26.2%) were found only in the water column (Figure 1B). In addition, very few ASVs were shared by both fractions (31 ASVs representing 6.7% of the total ASVs). This distribution of ASVs illustrates the statistical differences found in the beta-diversity analyses, highlighting a higher richness in the sediments than in the water samples. Consistent with the differences in beta diversity observed between the two different habitats (water column versus sediments), the distribution of two abundant Amoebozoa families appeared particularly contrasted between water samples and sediments. ASVs belonging to Paramoebidae were found specifically enriched in the sediments, whereas ASVs belonging to Vannellidae were found specifically enriched in the water column (ANCOM analyses; Table S4), which was not the case for the other Amoebozoa families, which tended to be more evenly distributed between the two habitats regardless of the sampling site.

**Figure 1.**
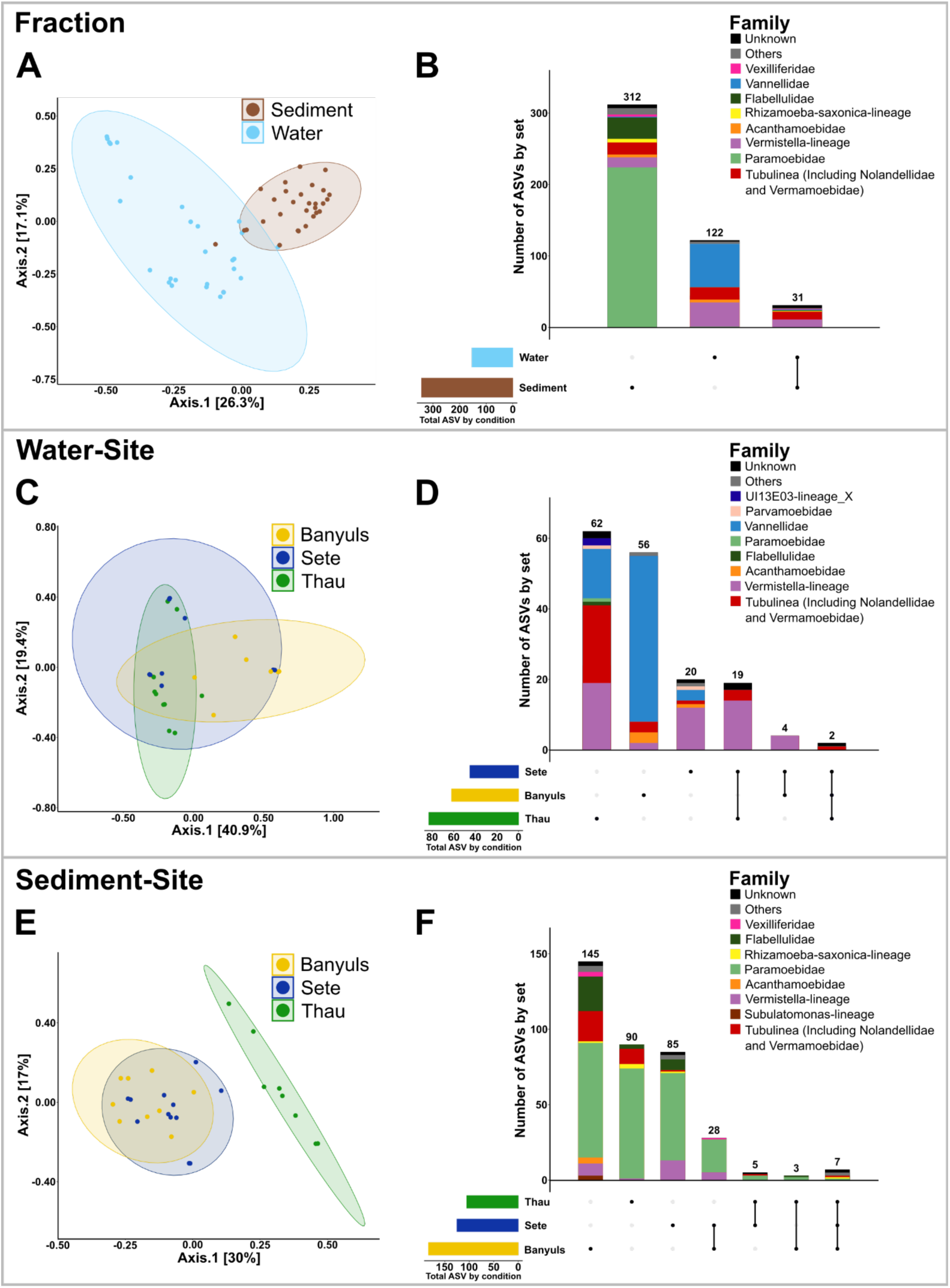
Amoebozoa diversity is structured according to sampling fraction and sampling site. PCoA of the unweighted Unifrac matrix distance revealed that the beta diversity of Amoebozoa in the sediment is different from the beta diversity in the water samples (A). The distribution of ASVs showed that ASVs belonging to Paramoebidae are particularly enriched in the sediment, whereas ASVs belonging to Vannellidae are particularly enriched in the water samples (B). PCoA of the unweighted Unifrac matrix distance between water samples from the different sampling sites showed that the beta-diversity of Amoebozoa in water is highly variable and does not differ significantly between sampling sites (C). ASVs distribution shows that most ASVs found in water samples are specific to each site and the community structure is variable, but ASVs of each family are found in all sites (D). The PCoA of the unweighted Unifrac matrix distance between the sediment samples of the different sampling sites showed that the beta diversity of Amoebozoa in the sediments is more homogeneous and the beta diversity found in the Thau sediments is different from the beta diversity in the sediments of Sète and Banyuls, which are more similar (E). The distribution of ASVs shows that most ASVs found in the sediments are specific to each site and the community structure is variable, but ASVs of each family are found in all sites (D).

### The diversity of cultivable marine Amoebozoa depends on the geographical location

Secondly, we investigated the differences in Amoebozoa diversity between the three contrasting sampling sites Banyuls-sur-Mer, Sète and Thau. No significant differences in alpha diversity were observed between the sites, but beta diversity using PCoA with the DEseq2 normalization method and Unifrac distance matrix showed that the beta diversity of each site was specific and significantly different from the others (Table S3). Due to the strong fraction effect, the taxonomic level of beta-diversity differences between sites was difficult to evaluate independently of fraction. Therefore, we compared the beta-diversity of Amoebozoa between sites in the water samples on the one hand and in the sediments on the other hand. Although the communities were all significantly different (Table S3), the three groups of samples overlapped (Figure 1C and 1D). These results suggest that the variance between samples belonging to the same site was large and in a similar range for each site. In fact, the community composition between the three sites was similar with intra-family abundance variation (Figure 1D). The repartition of ASVs by sites in the water column showed that out of the 163 ASVs, 62 ASVs (38%) and 56 ASVs (34.4%) were specific to Thau and Banyuls-sur-Mer, respectively, while only 20 ASVs (12.3%) were specific to Sète (Figure 1D). For the sediment samples, PCoA and Unifrac distance matrix showed that the beta diversity was significantly different between each site, with a little more homogeneity between samples from the same site than what was observed for the water samples (Figure 1E and 1F). The group of samples from Banyuls-sur-Mer and Sète overlapped, while the group of samples from Thau was very distinct, suggesting that the Amoebozoa communities in the sediments from Banyuls-sur-Mer and Sète are more similar between them than the community from Thau. The repartition of ASVs by site in the sediments showed that out of the 363 ASVs, 145 ASVs (39.9%) were specific to Banyuls-sur-Mer, while only 90 ASVs (24.8%) and 85 ASVs (23.4%) were specific to Thau and Sète, respectively (Figure 1F). Taken together, these results indicate that Amoebozoa beta-diversity is not stable and that different communities are observed between contrasting sites in the water column and in sediments. While the communities appear more stable in the sediments than in the water fraction, the Amoebozoa community assemblages are more specific to each site in the sediments. Overall, these results suggest that most of the taxonomic diversity specific to each environment appears to be at the subfamily taxonomic level, as a large majority of ASVs belonging to each family were found in all sampling sites, whereas a very limited number of ASVs (only 9 ASVs) belonging to different families were ubiquitous and common to all sampling sites. This was best illustrated by a phylogenetic analysis of all ASVs, which highlighted intra-family branches of ASVs sharing similar ecological characteristics such as sampling sites and fraction (Figure 3).

### Seasonal variations in Amoebozoa diversity occur at the subgenus level

We next attempted to evaluate seasonal variations in Amoebozoa diversity with respect to both fractions and the three sites. No significant differences were found for either alpha or beta diversity (Table S2 and S3). The repartition of ASVs between seasons revealed some noticeable differences with a higher number of ASVs found in summer and spring and a lower number of ASVs found in winter and fall (Figure 2A and 2B). In fact, of the 465 ASVs, 146 ASVs (31.4%) and 116 ASVs (24.9%) were specific to summer and spring, respectively, while only 73 ASVs (15.7%) and 24 ASVs (5.2%) were specific to winter and fall, respectively (Figure 2B). The taxonomic affiliations of the specific ASVs identified during summer and spring are very similar with some differences in numbers. They include ASVs belonging to the Rhizamoeba-saxonica lineage, Vermistella lineage, Tubulinea, and the families Flabellulidae, Acanthamoebidae, Vannellidae, and Paramoebidae (Figure 2B). Representatives of most Amoebozoa genera were present in winter and fall as well as in summer and spring, but the overall diversity was less dominated by ASVs belonging to the Paramoebidae family than in spring and summer (Figure 2B). Such variation at the subgenus level was clearly evident in the phylogenetic classification of ASVs, with some intra-genus clades being particularly present at a particular season (Figure 3). For example, among Paramoebidae, ASVs belonging to clade 11 were found only in summer in Thau lagoon, whereas among Vannellidae, ASVs belonging to clade 2 were found in winter in Thau lagoon, and ASVs belonging to clades 1 and 3 were found in summer in Banyuls (Figure 3). All these results suggest the presence of contrasting seasonal variations at the subgenus level, but our data set and the lack of taxonomic information did not allow to highlight statistically significant seasonal variations of Amoebozoa diversity.

**Figure 2.**
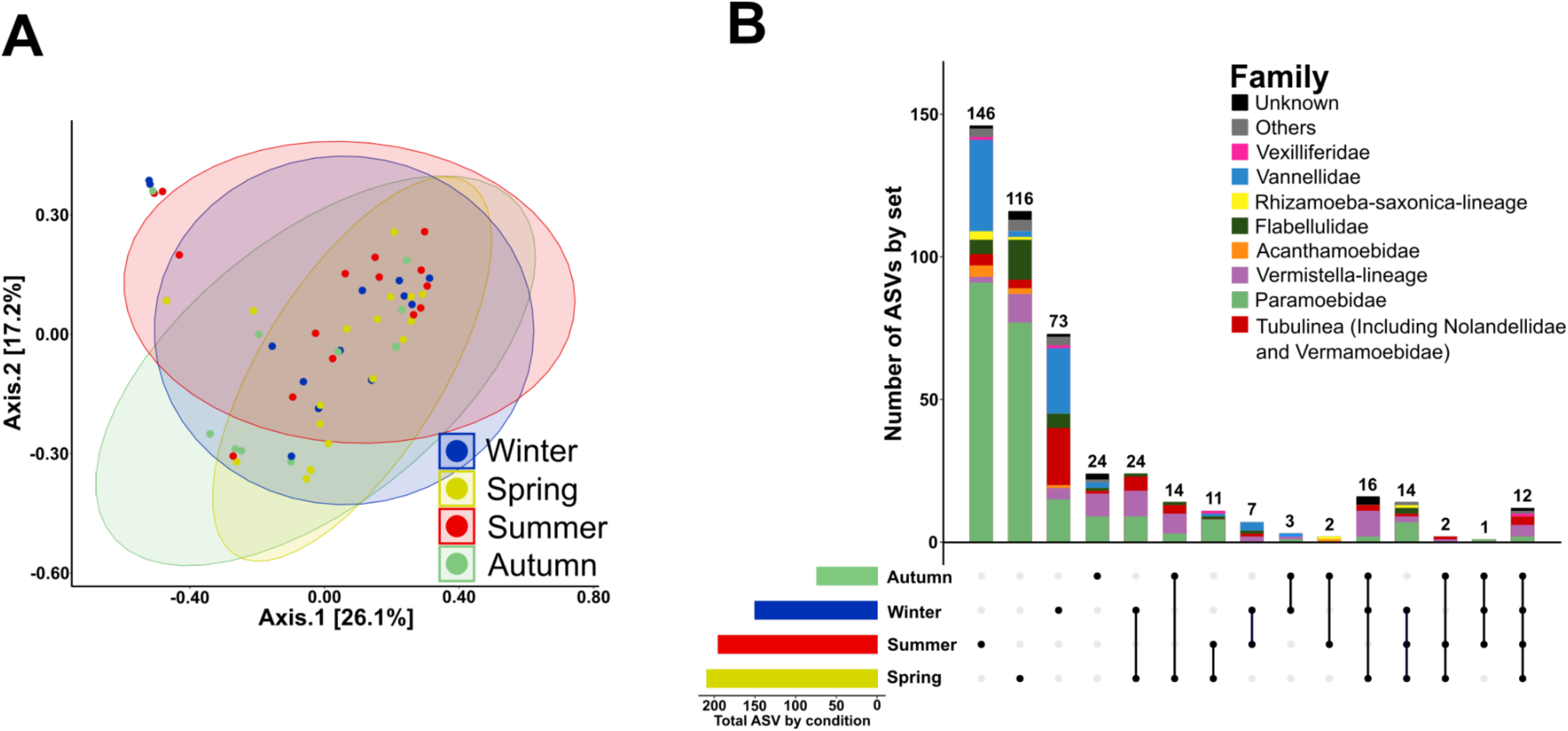
Seasonal variation of Amoebozoa diversity is very heterogeneous, but more ASVs are found in spring and summer than in winter and autumn. PCoA of unweighted Unifrac matrix distance revealed that the beta diversity of Amoebozoa found at different seasons is highly variable and overlaps between seasons (A). The distribution of ASVs showed that the total number of different ASVs is higher in spring and summer than in winter and autumn, ASVs belonging to each family are found in all seasons in different numbers (B).

**Figure 3.**
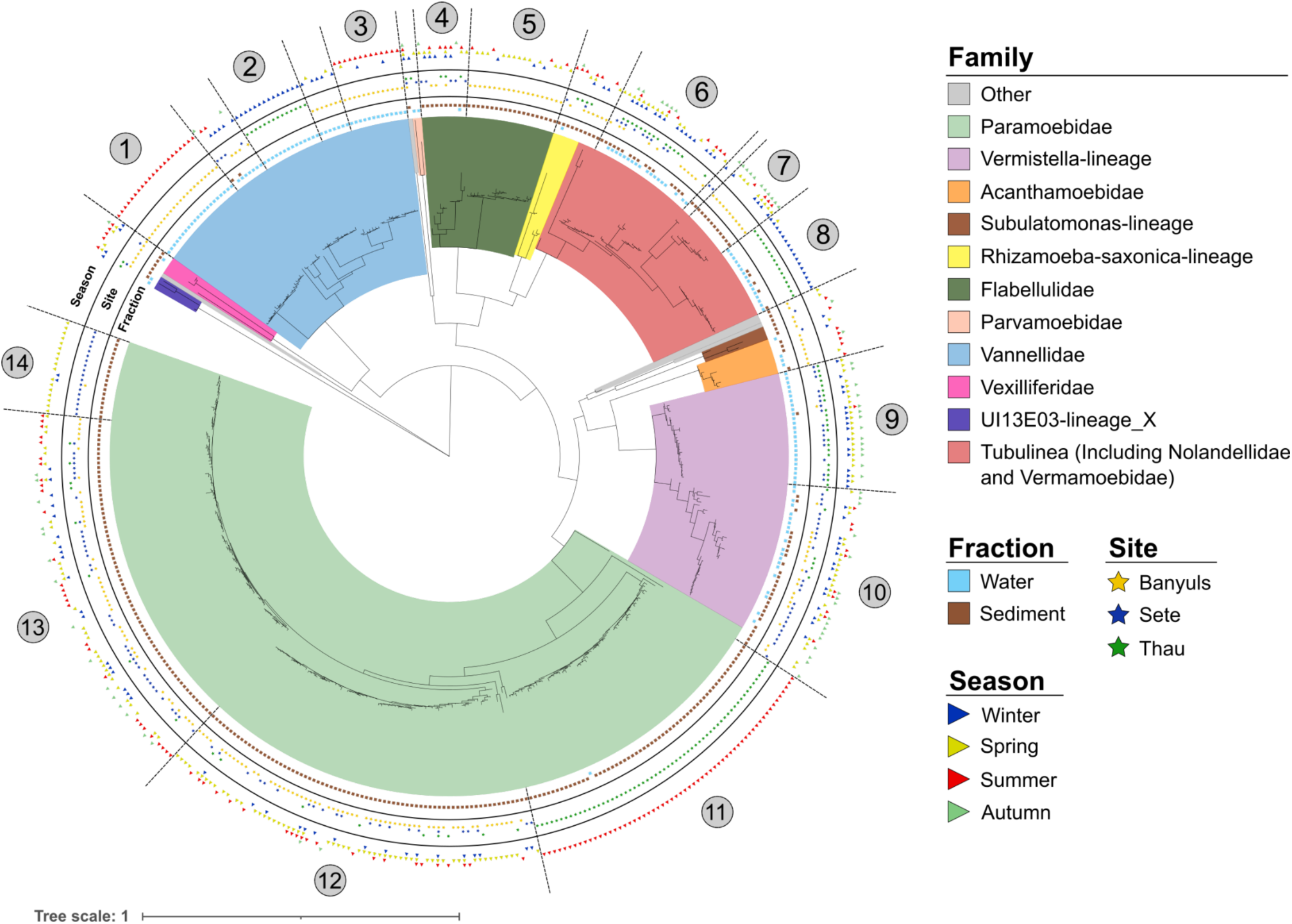
Phylogeny of ASVs highlighting intrafamilial clades of Amoebozoa present in different habitats. The phylogenetic classification of ASVs was performed using MAFFT and FastTree (maximum likelihood tree) and annotated using iTOL software, highlighting fraction, location, and season variables.

### Different Amoebozoa taxonomic groups have varying predation capacities against different pathogenic vibrios

Because the diversity of amoeba-vibrio interactions and their specificity remain poorly studied. We wondered whether different pathogenic vibrios with different anti-eucaryotic virulence mechanisms would select for different predators. Therefore, during four different months of the sampling campaign, *V. tasmaniensis* LGP32, *V. crassostreae* J2-9, and *V. harveyi* A01 were used as food sources in addition to the regular *E. coli* SBS363 (Figure S1). Since the abundance of pathogenic vibrios in the two different fractions, water column and sediments, differs during oyster mortality (Lopez-Joven et al., 2018), we chose 2 months during oyster mortality events (May 2017 and October 2017) and 2 months without mortality events (January 2018 and February 2018). The highest contrast in alpha diversity of Amoebozoa was observed between *V. harveyi* A01 plates and *V. tasmaniensis* LGP32 plates, as shown by all alpha diversity indices (Table S6), while *E. coli* SBS363 and *V. crassostrea*e J2-9 plates showed intermediate alpha diversity. Beta diversity didn’t show significant differences between the four prey species (Figure 4A and Table S6). The repartition of ASVs confirmed the results of alpha and beta diversity. Indeed, we identified the maximum number of specific ASVs and the maximum diversity with A01 plates (140 ASVs representing 29.5% of the 474 total ASVs), with specific ASVs belonging to the Paramoebidae family dominating the observed diversity and contrasting between the four different prey (Figure 4B). Of all detected ASVs, 58.7% were found on only one type of bacterial lawn, while 41.3% were found on at least two different types of bacterial lawn. Interestingly, 39 ASVs, representing 8.2% of the total ASVs and belonging to different Amoebozoa families, had the ability to feed and grow on the four different types of bacterial lawns, suggesting a less specific predatory activity (Figure 4B).

**Figure 4.**
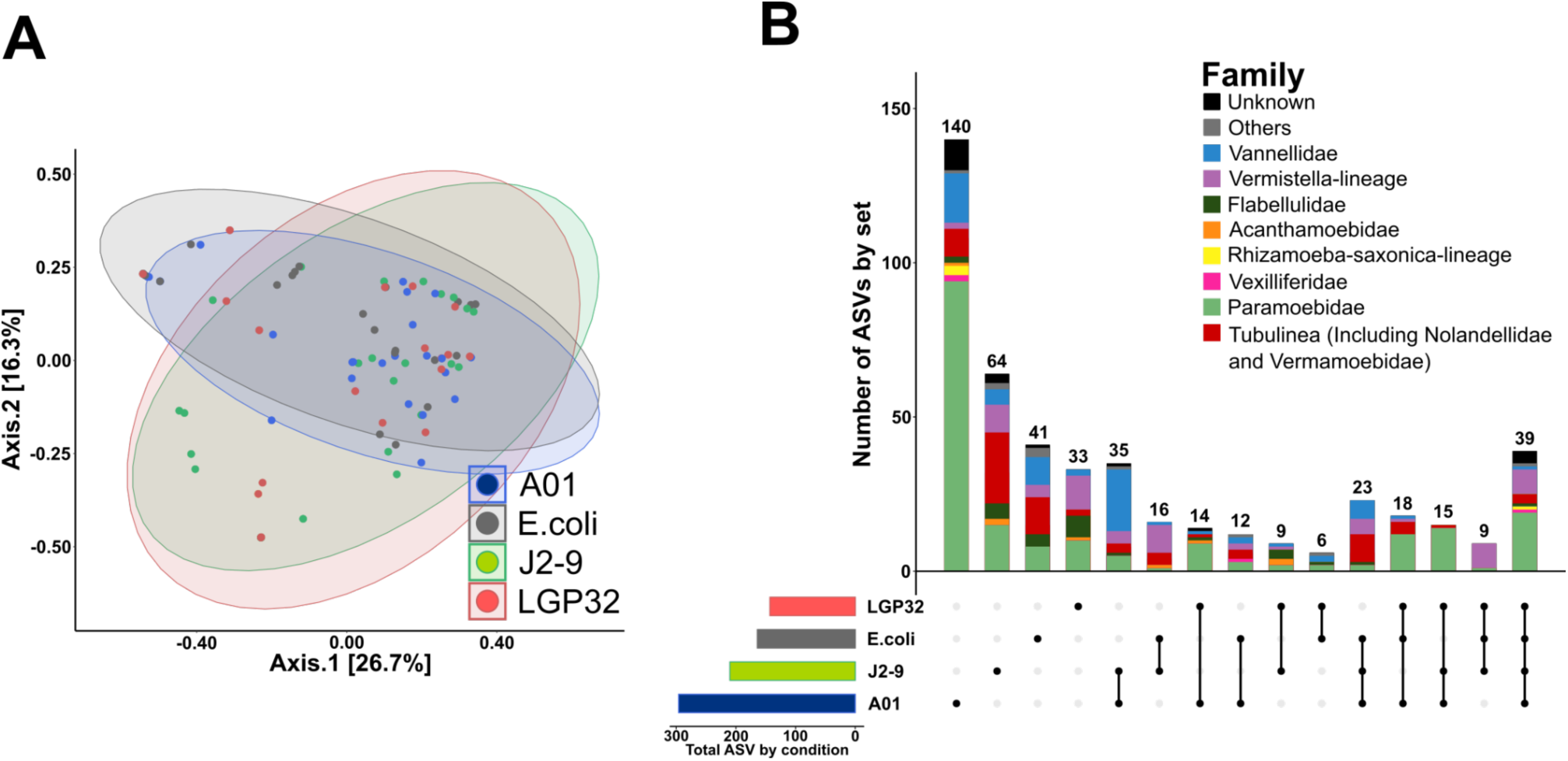
Selection on different bacterial prey revealed intra-family variation in the predatory capacity of amoebozoa against different opportunistic vibrios. PCoA of the unweighted Unifrac matrix distance revealed that the beta diversity of amoebozoa growing on the different bacterial prey is highly variable and mostly overlapping between conditions (A). The distribution of ASVs revealed that the total number of different ASVs is higher on *V. harveyi* A01 compared to the three other bacterial lawns, but 39 ASVs belonging to the different amoebozoa families were able to grow on the four different bacterial lawns (B).

### *Vibrio tasmaniensis* LGP32 strongly inhibited the growth of Vannellidae

Since the two most contrasting families appeared to be the Paramoebidae and Vannellidae in terms of sampling fraction, sampling sites, and predatory capacity, we analyzed these two families in more detail. Phylogenetic classification of Vannellidae ASVs revealed three distinct clades with different characteristics. Clade 1 was composed of relatively distant ASVs identified on the four bacterial lawns, mainly from the water samples with three ASVs from sediments at the three sites during the four seasons (Figure 5). Clade 2 gathers more closely related ASVs found mainly on the SBS363 lawn with 2 ASVs and 4 ASVs from LGP32 and A01 lawns, only in the water column, at Thau and during February (Figure 5). Clade 3 is composed of ASVs found mainly on A01 and J2-9 lawns with some ASVs found on SBS363 lawn, only in the water column and at Banyuls-sur-Mer and mainly during January and February with 2 ASVs identified during May (Figure 5). These observations illustrate well the existence of geographical and seasonal specificities as a matter of phylogenetic diversity of Vannellidae. In addition, amoebae belonging to the Vannellidae appeared to be much less prone to grow on the LGP32 lawn than on the other bacterial lawns; only a few ASVs were identified from samples grown on the LGP32 lawn. These results are reminiscent of our previous study showing that LGP32 is resistant to predation by the amoeba *Vannella* sp. AP1411 (Robino et al., 2020).

**Figure 5.**
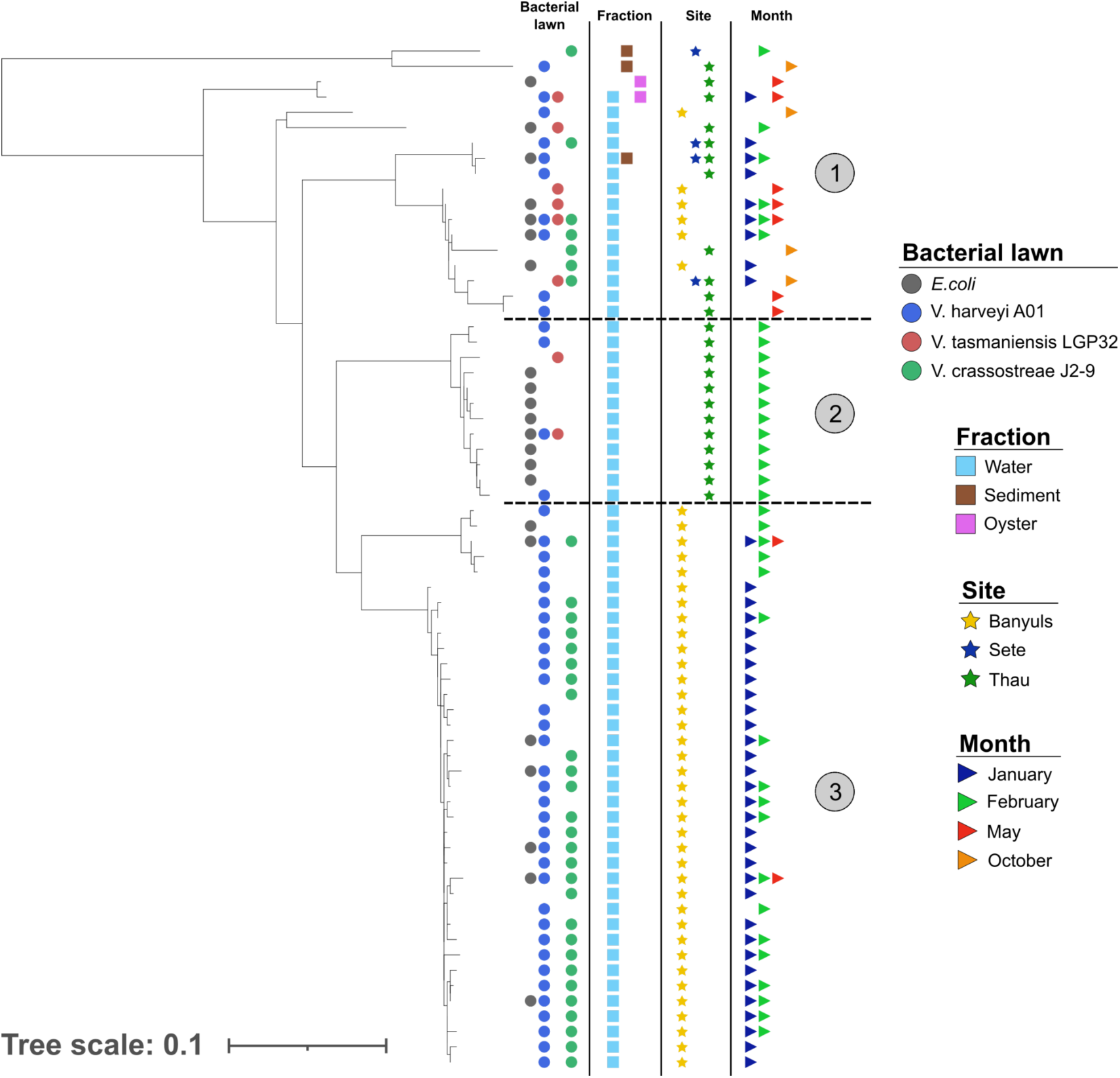
Phylogeny of ASVs highlighting clades of Vannellidae found in different environments and with different predatory capacities. The phylogenetic classification of ASVs was performed using FastTree (maximum likelihood tree) and annotated using iTOL software, highlighting fraction, location, and season variables. Clade 1 was heterogeneous and found in different conditions but was able to grow on any of the bacterial prey, clade 2 was found in Thau in February and grew mainly on E.coli, and clade 3 was found mainly in Banyuls in January and February and was unable to grow on LGP32.

### The Paramoebidae represent a highly diversified taxonomic group with diverse predatory capabilities

The Paramoebidae represented the amoeboid family with the highest number of identified ASVs (Figure 6). Furthermore, this family appeared to contain ASVs with some having a tendency towards generalist predatory capacity and some others appearing to be more specialized, but most of them showed a high capacity to feed on *V. harveyi* A01. There were two clades (clade 3 and 5) able to feed on the four food sources, mainly found in the sediments, at the three sites for clade 3 with a majority at Banyuls-sur-Mer and Sète for clade 5, during the four months sampled with fewer ASVs in October. In addition, four more specialized clades (clades 1, 2, 4 and 6) were identified, showing some differences. Clades 1 and 4 were found similarly in the water column, at Banyuls-sur-Mer and in October. These two clades are interesting because very few of the total ASVs were identified during October. Clade 2 and 6 are similar and were mainly identified at Sète and during May. However, ASVs of the clade 2 were mainly found in the water column and clade 6 in the sediments. Thus, these results highlight that the Paramoebidae were the largest family of Amoebozoa sampled here and that their ecological dynamics as well as their predatory capacities can vary greatly.

**Figure 6.**
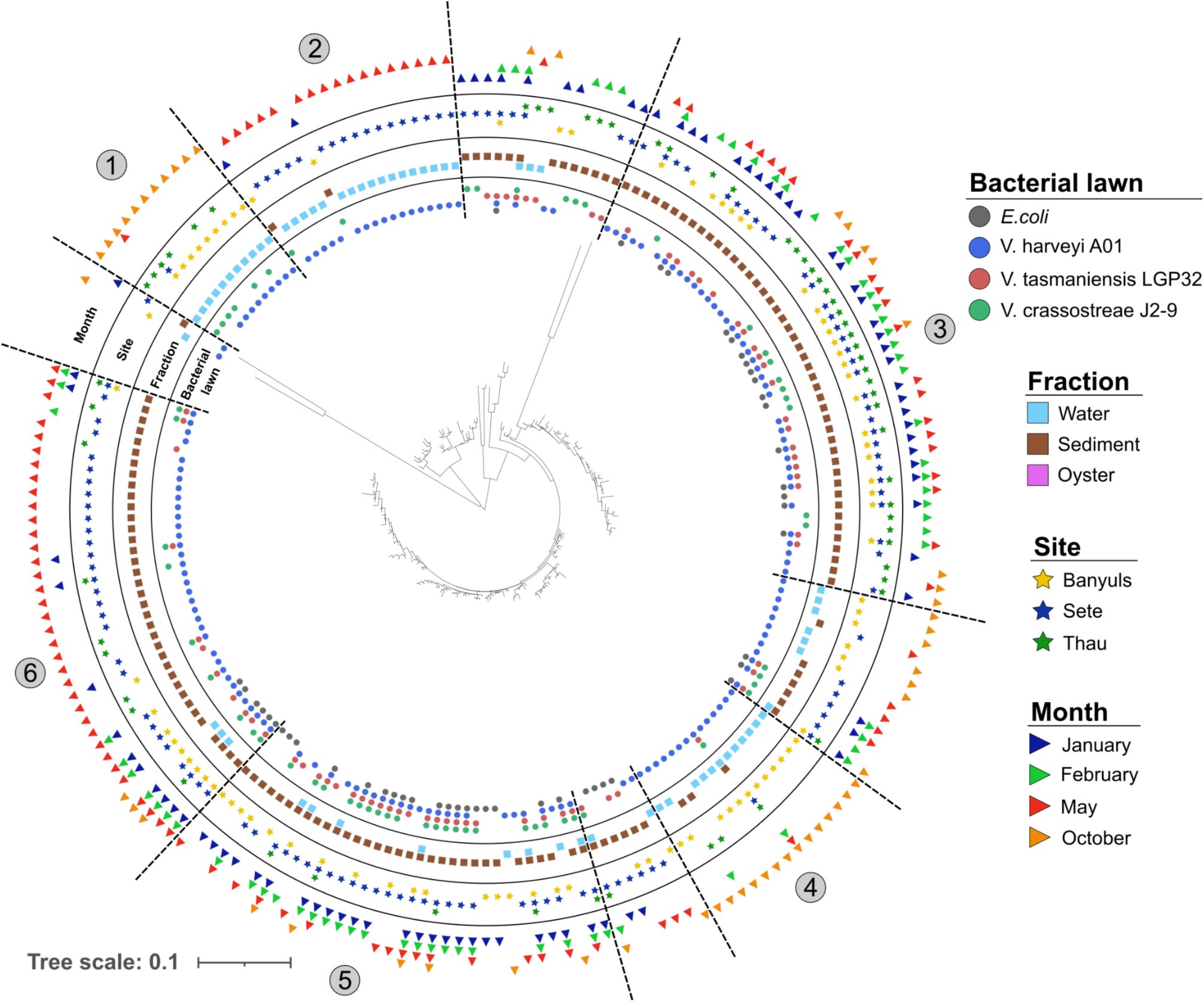
Phylogeny of ASVs highlighting different clades of Paramoebidae found in different environments with different predation capacities. The phylogenetic classification of ASVs was performed using MAFFT and FastTREE FastTree (maximum likelihood tree) and annotated using iTOL software, highlighting fraction, site, and season variables. Some clades appeared to be ubiquitous and mostly generalist as they could grow on the four bacterial lawns, such as clade 3 and 5, whereas other clades appeared to have more restricted habitats with a more limited predation capacity.

### Contrasting predation capacities against phylogenetically related vibrios of the Splendidus clade were functionally validated using Vannellidae and Paramoebidae isolates

To further confirm that Vannellidae growth was inhibited by *V. tasmaniensis* LGP32 and Paramoebidae growth was inhibited by *V. crassostrae* J2-9, we first compared the total abundance of ASVs belonging to these two families among all the samples obtained on LGP32 and J2-9 lawns. The relative abundance of Paramoebidae was found to be significantly higher than that of Vannellidae on LGP32 lawn (Fisher test p<0.05; Figure 7A). On the contrary, the abundance of ASVs belonging to Paramoebidae on J2-9 lawn was found to be significantly lower than the abundance of ASVs belonging to Vannellidae (Fisher Test p<0.05). These results suggest that LGP32 inhibits the growth of Vannellidae while Paramoebidae can feed on this vibrio strain, and on the contrary, J2-9 tends to inhibit the growth of Paramoebidae while Vannellidae grows better using this strain as a food source. To functionally validate these results, we performed grazing experiments with LGP32-GFP and J2-9-GFP strains to quantify the predation activity of *Vannella sp.* AP1411 and *Paramoeba atlantica* (strain CCAP1560/9) over time. The results showed that *Vannella sp.* AP1411 could not graze on LGP32 (as previously published by Robino *et al*. 2019), whereas J2-9 was rapidly eliminated, as shown by the rapid decrease in the relative fluorescence of GFP, which was reduced to a ratio of 0.4 after five days (ANOVA, p<0.001; Figure 7B). In contrast, grazing experiments with *Paramoeba atlantica* showed that LGP32 was grazed faster than J2-9, although the differences between the two vibrios strains were less pronounced than *Vannella sp.* AP1411 (ANOVA, p<0.05; Figure 7C). Taken together, these functional data confirmed that the two genera of Vannellidae and Paramoebidae may have different predatory capacities against different pathogenic vibrios belonging to the Splendidus clade

**Figure 7.**
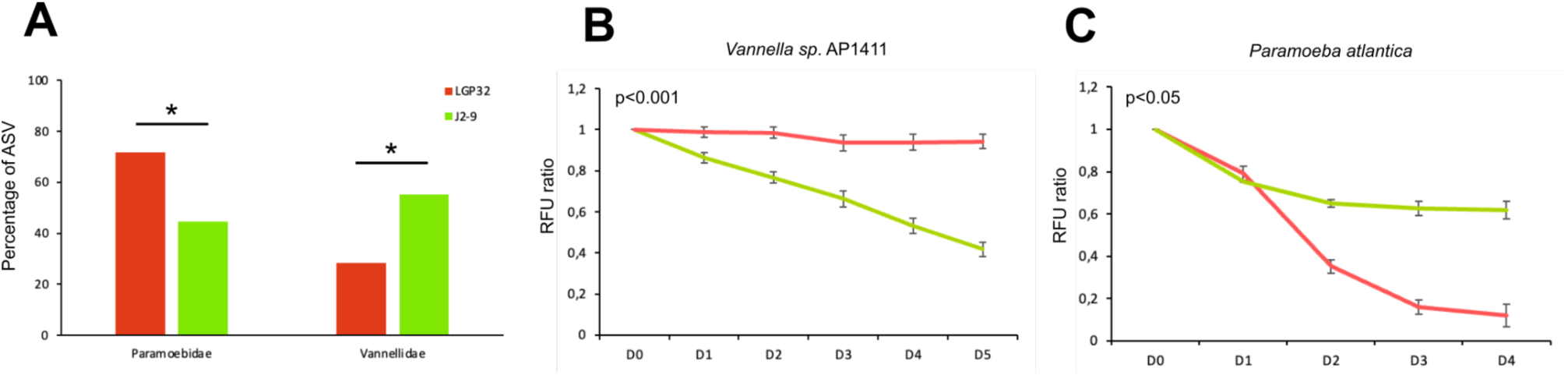
*V. tasmaniensis* LGP32 and *V. crassostreae* J2-9 show contrasting resistance to predation by Vannellidae and Paramoebidae. <The abundance of ASVs belonging to Paramoebidae or Vannellidae growing on LGP32 or J2-9 turf was significantly different (Fisher test; p<0.05) (A). Grazing experiments with *Vannella sp.* AP1411 showed that J2-9 was more sensitive than LGP32 (one experiment representative of three independent experiments, RM-ANOVA test; p<0.001). Grazing experiments with *Paramoeba atlantica* showed that J2-9 was more resistant than LGP32 (one experiment representative of three independent experiments, RM-ANOVA test; p<0.05).

## Discussion

To investigate the ecology of FLA populations and their interactions with *Vibrionaceae* in the Mediterranean coastal environment, we conducted monthly sampling for one year in three contrasting habitats. Free-living amoeba populations were isolated by culturing water and sediment samples on different bacterial lawns, including *E. coli* SBS363 or *V. tasmaniensis* LGP32, *V. crassostreae* J2-9 and *V. harveyi* A01. Analysis of protist diversity in the different samples by v4-18S rRNA barcoding revealed distinct communities of Amoebozoa between the sediments and the water column, with Vannellidae significantly enriched in the water column and Paramoebidae significantly enriched in the sediments. Moreover, the diversity of Amoebozoa in the sediments was more specific to the sampling sites than in the water column. Selection of grazers on different bacterial lawns revealed that *V. tasmaniensis* LGP32 inhibited the growth of most Vannellidae, whereas *V. crassostreae* J2-9 tended to inhibit the growth of Paramoebidae. These differences were further confirmed in functional grazing assays using isolates belonging to each Amoebozoa taxonomic group. Overall, our results highlight the need for more comprehensive studies of amoeboe diversity and population dynamics in marine waters, and the role of these diversified grazers in shaping vibrio communities and pathogen dynamics remains poorly characterized.

In contrast to our previous observations, by conducting a more extensive field survey, we found, in addition to Vannellidae, many other Amoebozoa belonging to Paramoebidae, Tubulinea (regrouping Vermamoebidae and Nollandelidae), Rhizamoeba-saxonica, Vermistellae, Flabellulidae, Vexilliferidae, and Parvamoebidae families. Although some were more abundant than others, including Paramoebidae, Vannellidae, Vermistellae, Tubulineae, and Flabellulidae, most had been previously reported from various marine environments (Garstecki and Arndt, 2000; Latifi et al, 2020; Mohd Hussain et al., 2022; Munson, 1992; Page, 1983; Samba-Louaka et al., 2019; Sousa-Ramos et al., 2022). A limitation of our study was the use of universal v4-18S primers to analyze FLA communities. FLA are very diverse and paraphyletic and the use of universal v4-18S primers may have precluded to identify the full diversity, especially in taxonomic phylum other than Amoebozoa such as Heterolobosea for example (Delafont et al., 2022). Besides Amoebozoa, the other abundant phylum of grazers detected here was Cercozoa, but the taxonomic assignment was limited due to the lack of reference sequences in 18S databases. Even for Amoebozoa, taxonomic assignment based on 18S rRNA partial sequences can be inconsistent at the species level and in some cases even at the genus level, which is the reason why we analyzed the diversity of Amoebozoa mostly at the family level.

A limited number of studies have performed a systematic sampling survey over time to compare the diversity of FLA between sites and between different fractions in marine environments as reported here. Overall, the higher diversity of Amoebozoa observed in sediments compared to water samples is consistent with their benthic grazer lifestyle. Distinct grazer communities between sediments and water column have also been reported by other studies that found different protist communities between surface and deep waters of the same site as well as between different sediment depths (Countway et al., 2007; Orsi et al., 2011; Smirnov, 2004). Here, several ASVs belonging to the different families were found in the different environments, suggesting that most of the diversity differences appear to be at the intrafamily and probably intrageneric level. Nevertheless, a limited number of ASVs were found in all sampling sites, suggesting that cosmopolitan strains of Amoebozoa are rare. However, the distribution of ASVs between the two different habitats was particularly contrasting for the Vannelliadae, which were specifically enriched in the water column, and the Paramoebidae, which were specifically enriched in the sediments. This may be related to a specific lifestyle and adaptations, as Vannellidae are known to form characteristic elongated filopodia in their planktonic form, which may provide an advantage for swimming and movement using water currents (Smirnov et al., 2007). In contrast, Paramoebidae are frequently reported in marine sediments and are rare in freshwaters, suggesting a specialized marine benthic lifestyle (Page, 1983; Kudryavtsev et al., 2011; Volkova et al., 2019; Volkova and Kudryavtsev, 2017). Amoebozoa communities in the sediments were found here to be more stable over the sampling months but contrasting between sites, whereas communities from the water samples were found to be more variable and less contrasted between sampling sites. This may be explained by differences in the physical characteristics and variability of the two fractions. The water column is subject to significant variability and rapid mixing by water movement and currents, increasing the connectivity between environments, in contrast to the sediments, which are more preserved from mixing events. In addition, the physicochemical composition of sediments between sites (e.g., depth, sand/mud composition, and oxygen concentrations) is likely to be different, which could have a strong influence on the niche characteristics and composition of microorganism communities (Kim et al., 2014; Smirnov, 2004). Further efforts in exhaustive comparative analyses between contrasting environments are still needed to unravel the most important environmental factors shaping FLA communities.

Unfortunately, the analysis of seasonal variation was hampered by the heterogeneity between samples over the sampling period. Nevertheless, the distribution of ASVs between seasons showed that the number of different ASVs was higher in spring and summer and lower in fall and winter (Figure 4B). These observed trends are consistent with other studies showing seasonality in protist diversity (Berdjeb et al., 2018; Fu et al., 2020; Kim et al., 2014). Phylogenetic analysis revealed that within each family, some clades of ASVs appeared to be either ubiquitous and found in all fractions, all sites, and all seasons, while other clades of ASVs appeared to be more restricted. The Tubulinea taxon is the best example to illustrate this, as it contains three completely different taxonomic groups of ASVs (Figure S6). One of them tends to be ubiquitous, found in both fractions, at the 3 sites and during the four seasons (clade 6). The other two clades (clades 7 and 8) appear to be more specialized, showing several differences that suggest two different lifestyles. A more resolutive sampling strategy may be required to fully capture the seasonal variation of the different taxonomic groups.

Biotic factors such as prey can also influence the composition of heterotrophic protist communities. Some prey have evolved the ability to resist predation using various extracellular and intracellular mechanisms, sometimes even killing them (Matz and Kjelleberg, n.d.; Pernthaler, 2005; Robino et al., 2020). Amaro and his collaborators have shown that *Legionella pneumophila* can shape protist communities in microcosm experiments, with significant effects on the abundance of the phyla Cercozoa, Amoebozoa, and Heterolobosea (Amaro et al., 2015). Here, we wondered whether three different strains of opportunistic vibrios with different anti-eukaryotic virulence mechanisms could influence the diversity of FLA communities. Surprisingly, the highest diversity was observed when *V. harveyi* A01 was used as a food source compared to the other strains, which showed three times more specific ASVs than *E. coli* SBS363 (Figure 4B). On the contrary, *V. tasmaniensis* LGP32 and to a lesser extent *V. crassostreae* J2-9 were the most selective bacterial lawns as we found a lower amount of specific ASVs, suggesting that their anti-eucaryote defense mechanisms could be efficient against a greater diversity of amoebozoa. We observed more generalist or more specialist clades of ASVs in most Amoebozoa families, with some ASVs that could grow with any of the prey, suggesting the existence of generalist species with a wide range of prey and habitats. However, their number was limited and many ASVs had more specialized grazing capacity towards the different prey. Among them, the majority of Vannellidae and a large part of Tubulinea could not grow in the presence of LGP32. This is reminiscent of our previous study showing that LGP32 is resistant to predation by *Vannella sp*. AP1411 (Robino et al., 2020). Paramoebidae was the family that contained the highest number of different ASVs, but their growth was particularly inhibited by J2-9. The contrasts in prey specificity of Vannellidae and Paramoebidae were functionally confirmed using *Vannella sp.* AP1411 and *Paramoeba atlantica* with both vibrios. These results emphasize that although intrageneric variations in predation capacity are observed, the resistance to grazing of some vibrios, such as LGP32 and J2-9, can affect larger taxonomic groups of Amoebozoa. Seasonal variations in the abundance of *V. tasmaniensis* and *V. crassostreae* in different habitats have been previously reported in the Thau lagoon (Lopez-Joven et al., 2018). These vibrios were found more abundant in the water column during the warmer seasons, but restricted to the sediment during the colder seasons. Because Vannellidae are more abundant in the water column and Paramoebidae are more abundant in the sediment, the differences in Amoebozoa diversity in the two different habitats could have opposite effects in shaping vibrio communities and differentially affect the dynamics of different pathogens over the seasons. Similarly, *in vitro* functional grazing assays have been used by others to highlight differential predation capacity and prey specificity and to predict potential effects on bacterial communities in the plant rhizosphere (Amacker et al., 2022). Interestingly, some interactions between vibrios and Paramoebidae have been reported previously; their role in vibrio population dynamics and as a potential intracellular niche needs further investigations (Lee et al., 2013; MacPhail et al., 2021). .

In conclusion, our study provides a better understanding of Amoebozoa grazer communities in Mediterranean coastal environments and sets the stage for further in-depth functional studies between different Amoebozoa taxonomic groups and the bacterial communities present in these environments. By using different marine vibrios, we bring new evidence that Amoebozoa-vibrio interactions are highly diverse and underline the need to further study Amoebozoa diversity and their role in vibrio community dynamics and pathogen emergence.

## Acknowledgements

We are grateful to Eve Toulza, Jérémie Vidal Dupiol, and Jean-Christophe Auguet for fruitful discussions and precious help in sequencing analysis. We thank Philippe Haffner and Marc Leroy for technical assistance. This work; through the use of the GENSEQ platform (http://www.labex-cemeb.org/fr/genomique-environnementale-2) from the labEx CeMEB. The present study was supported by the Ec2co-CNRS funded VibrAm project, by the UE funded project VIVALDI (H2020 program, No. 678589), by the EU funded EMBRC and by Ifremer, University of Montpellier and University of Perpignan via Domitia.

## Supplementary Figures

**Figure S1.**
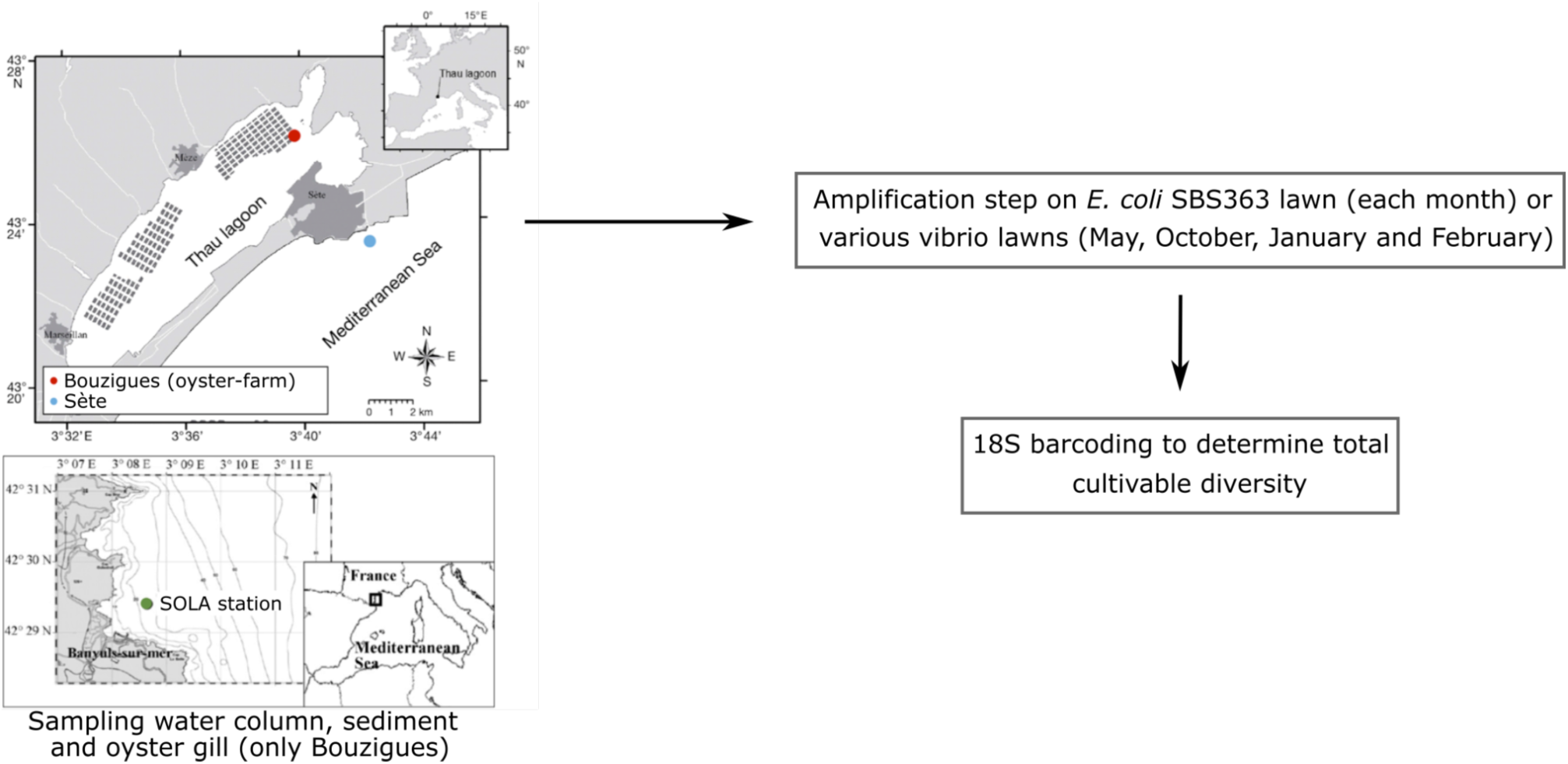
Simplified diagram of the sampling strategy. Briefly, water column and sediment were sampled monthly for one year in the lagoons of Sète, Banyuls-sur-Mer and Thau, near the Bouzigues oyster farming area. Grazers were isolated by selective growth and migration on agar plates coated with E. coli SBS363 lawns. In May, October, January and February, grazer isolation from the same samples was additionally performed on vibrios lawns (V. harveyi A01, V. tasmaniensis LGP32 and V. crassostreae J2-9). All grazers that grew and migrated from the initial sample were recovered and total DNA was extracted for v4-18S barcoding.

**Figure S2.**
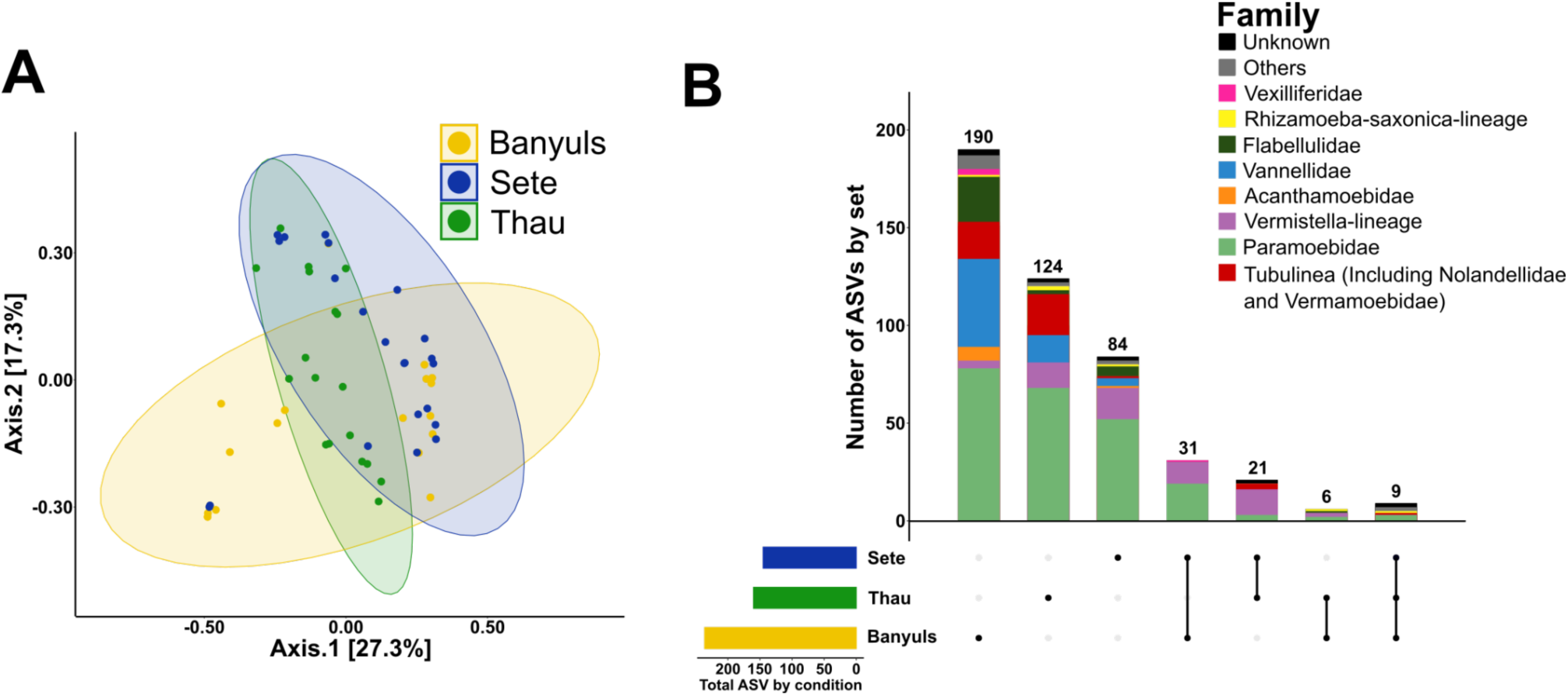
Amoebozoa diversity according to sampling site. PCoA of the unweighted Unifrac matrix distance showing the beta diversity of Amoebozoa in all samples and fractions between the three sampling sites (A). The distribution of ASVs showed that more ASVs were identified in Banyuls-sur-Mer compared to the other two sites, very few ASVs were present in more than one site, and only 9 ASVs were found in all sites (B).

**Figure S3.**
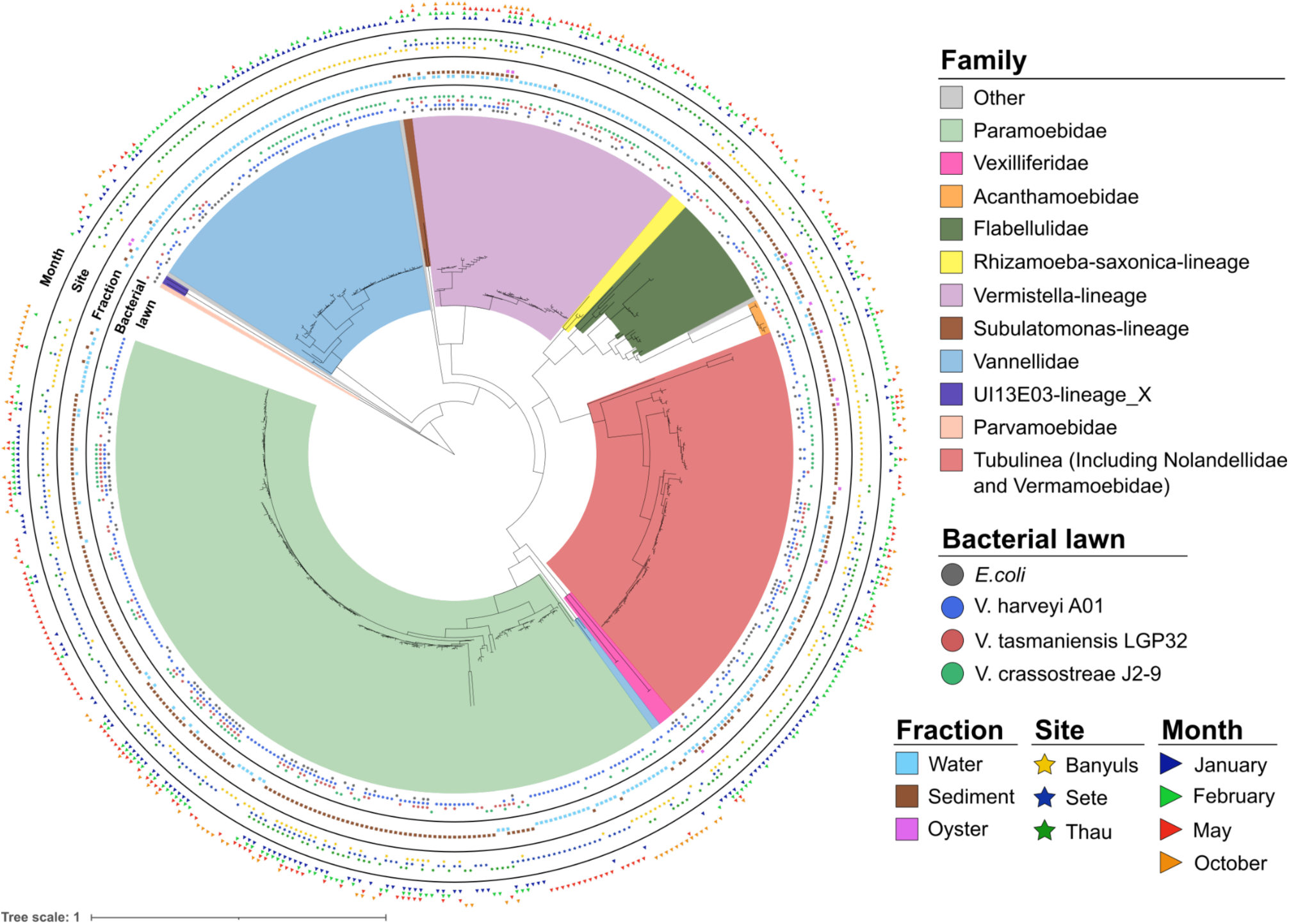
Phylogeny of all ASVs identified on the four different bacterial lawns. The phylogenetic classification of the ASVs was performed using MAFFT and FastTREE FastTree (Maximum Likelihood tree) and annotated using iTOL software, highlighting fraction, location, and season variables.

**Figure S4.**
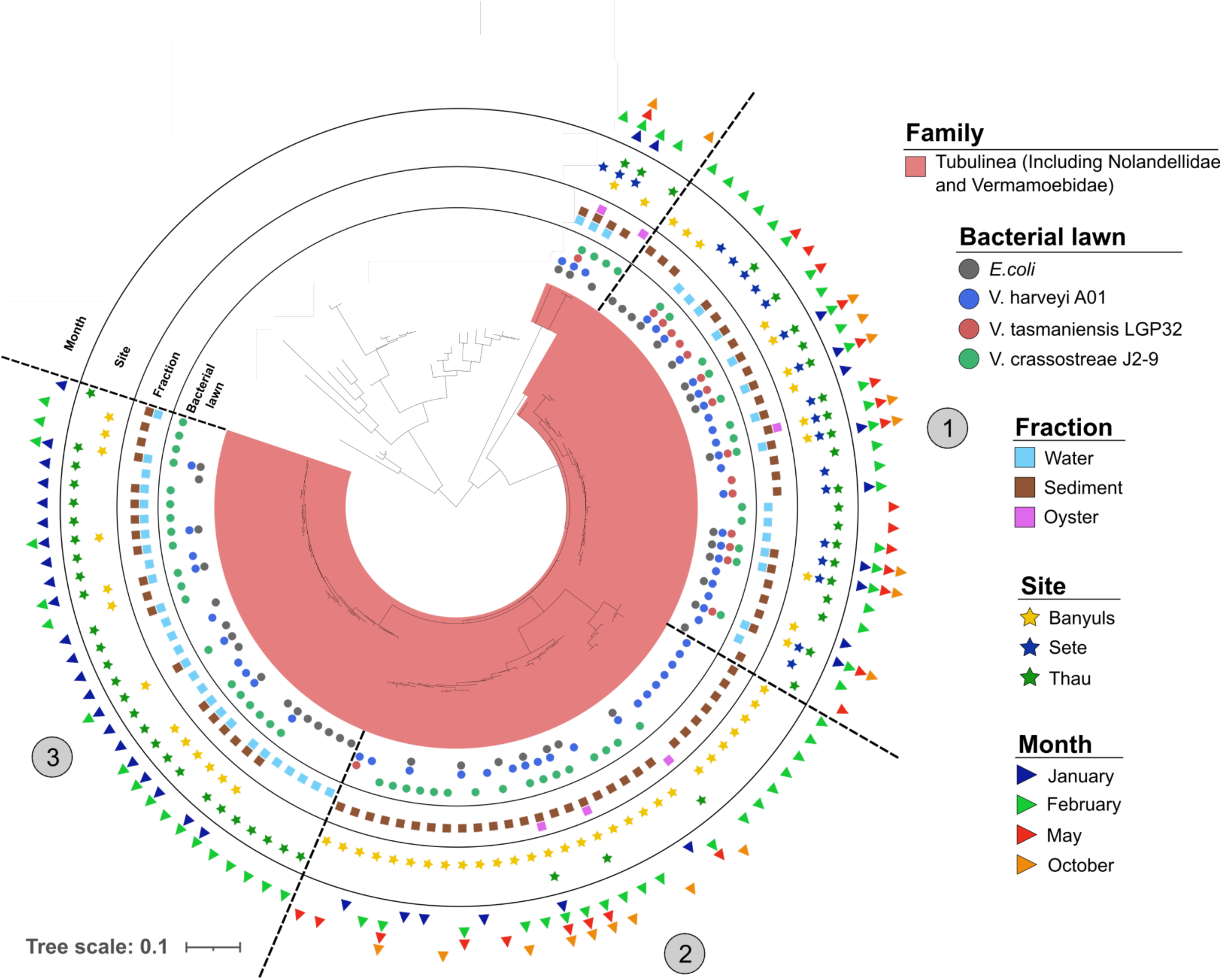
Phylogeny of ASVs highlighting different clades of Tubulinea found in different environments with different predation capacities. The phylogenetic classification of ASVs was performed using MAFFT and FastTREE FastTree (Maximum Likelihood tree) and annotated using iTOL software, highlighting fraction, location and season variables. Some clades appeared to be ubiquitous and mostly generalist as they could grow on the four bacterial lawns like clade 1, whereas clades 2 and 3 appeared to have more restricted habitats with a more limited predation capacity.

**Table S2.** (A) Statistics of the alpha diversity using the Chao1, Shannon and InvSimpson diversity indexes on Fraction, Site and Season variables. ANOVA on Chao1, Shannon and InvSimpson diversity indexes

|  |  | Chao1 |  | Shannon |  | InvSimpson |  |
| --- | --- | --- | --- | --- | --- | --- | --- |
|  |  | F value | Pr (>F) | F value | Pr (>F) | F value | Pr (>F) |
| **Fraction** | Water column vs Sediment | 6.525 | 0.0138 (*) | 3.734 | 0.0591 | 2.349 | 0.132 |
| **Site** | Banyuls vs Sète | 0.698 | 0.40912 | 0.002 | 0.961 | 0.025 | 0.875 |
|  | Banyuls vs Thau | 0.978 | 0.330 | 0.010 | 0.922 | 0.967 | 0.333 |
|  | Sète vs Thau | 0.078 | 0.782 | 0.189 | 0.667 | 1.061 | 0.310 |
| **Season** | Winter vs Autumn | 1.027 | 0.32244 | 1.984 | 0.174 | 0.133 | 0.719 |
|  | Winter vs Spring | 0.114 | 0.738 | 0.113 | 0.7394 | 0.466 | 0.500 |
|  | Winter vs Summer | 0.383 | 0.5415 | 0.199 | 0.659 | 0.020 | 0.890 |
|  | Autumn vs Spring | 1.432 | 0.242 | 3.023 | 0.0939 | 0.712 | 0.406 |
|  | Autumn vs Summer | 0.551 | 0.466 | 0.097 | 0.75809 | 0.010 | 0.921 |
|  | Spring vs Summer | 0.000 | 0.990 | 0.599 | 0.445 | 0.231 | 0.634 |

|  |  | Chao1 |  | Shannon |  | InvSimpson |  |
| --- | --- | --- | --- | --- | --- | --- | --- |
|  |  | F value | Pr (>F) | F value | Pr (>F) | F value | Pr (>F) |
| **Water** | Banyuls vs Sète | 1.09 | 0.313 | 0.426 | 0.523895 | 0.469 | 0.504 |
|  | Banyuls vs Thau | 0.439 | 0.518 | 0.483 | 0.4977 | 1.562 | 0.2305 |
|  | Sète vs Thau | 2.807 | 0.112 | 1.413 | 0.25093 | 2.589 | 0.1260 |

|  |  | Chao1 |  | Shannon |  | InvSimpson |  |
| --- | --- | --- | --- | --- | --- | --- | --- |
|  |  | F value | Pr (>F) | F value | Pr (>F) | F value | Pr (>F) |
| **Sediment** | Banyuls vs Sète | 1.110 | 0.306 | 0.054 | 0.819 | 0.000 | 0.997 |
|  | Banyuls vs Thau | 1.770 | 0.205 | 0.325 | 0.5775 | 0.055 | 0.817 |
|  | Sète vs Thau | 0.023 | 0.881 | 0.122 | 0.73094 | 0.054 | 0.8182 |

**Table S3.**

|  |  | R<sup>2</sup> | Pr (>F) |
| --- | --- | --- | --- |
| **Fraction** | Water column vs Sediment | 0.19817 | 0.0001 (***) |
| **Site** | Banyuls vs Sète | 0.05936 | 0.0396 (*) |
|  | Banyuls vs Thau | 0.11076 | 0.0001 (***) |
|  | Sète vs Thau | 0.12165 | 0.0002 (***) |
| **Season** | Winter vs Autumn | 0.04621 | 0.385 |
|  | Winter vs Spring | 0.02893 | 0.5399 |
|  | Winter vs Summer | 0.02488 | 0.7264 |
|  | Autumn vs Spring | 0.04115 | 0.2992 |
|  | Autumn vs Summer | 0.04445 | 0.3417 |
|  | Spring vs Summer | 0.05616 | 0.065 |

|  |  | R<sup>2</sup> | Pr (>F) |
| --- | --- | --- | --- |
| **Water** | Banyuls vs Sète | 0.1262 | 0.0138 (*) |
|  | Banyuls vs Thau | 0.25348 | 0.0012 (**) |
|  | Sète vs Thau | 0.11507 | 0.0213 (*) |

|  |  | R<sup>2</sup> | Pr (>F) |
| --- | --- | --- | --- |
| **Sediment** | Banyuls vs Sète | 0.11439 | 0.0044 (**) |
|  | Banyuls vs Thau | 0.27517 | 0.0001 (***) |
|  | Sète vs Thau | 0.25265 | 0.0001 (***) |

**Table S6.** (A) Statistics of the alpha diversity using the Chao1, Shannon and InvSimpson diversity indexes on Strain variable. ANOVA on Chao1, Shannon and InvSimpson diversity index

|  |  | Chao1 |  | Shannon |  | InvSimpson |  |
| --- | --- | --- | --- | --- | --- | --- | --- |
|  |  | F value | Pr (>F) | F value | Pr (>F) | F value | Pr (>F) |
| Strain | SBS363 vs A01 | 4.964 | 0.0313 (*) | 1.063 | 0.308 | 2.056 | 0.159 |
|  | SBS363 vs LGP32 | 0.275 | 0.602953 | 3.293 | 0.0769 | 3.279 | 0.0775 |
|  | SBS363 vs J2-9 | 0.237 | 0.62847 | 0.176 | 0.677 | 0.036 | 0.850 |
|  | A01 vs LGP32 | 6.951 | 0.011692 (*) | 7.632 | 0.00847 (**) | 9.680 | 0.00334 (**) |
|  | A01 vs J2-9 | 2.646 | 0.11076 | 1.523 | 0.224 | 1.183 | 0.282 |
|  | LGP32 vs J2-9 | 0.792 | 0.378344 | 1.652 | 0.205 | 2.397 | 0.129 |

|  |  | R<sup>2</sup> | Pr (>F) |
| --- | --- | --- | --- |
| Strain | SBS363 vs A01 | 0.01926 | 0.5398 |
|  | SBS363 vs LGP32 | 0.02299 | 0.4333 |
|  | SBS363 vs J2-9 | 0.02015 | 0.481 |
|  | A01 vs LGP32 | 0.02948 | 0.22 |
|  | A01 vs J2-9 | 0.03109 | 0.1376 |
|  | LGP32 vs J2-9 | 0.0178 | 0.61 |

